# Enhancing Visuomotor Reaction during Dark Transition via Model-Optimized Transcutaneous Vagus Nerve Stimulation

**DOI:** 10.64898/2026.10.17.759798

**Authors:** Shintaro Nakatani, Wuyuqing Yang, Joshua N. Posen, Jun Ueda, Minoru Shinohara

## Abstract

Sudden reductions in ambient illuminance temporarily impair visuomotor performance due to the inherent latency of dark adaptation, posing significant risks in operational environments such as tunnel entry. To mitigate this degradation, we propose a model-based neuromodulation framework using transcutaneous vagus nerve stimulation (tVNS), which is known to accelerate and augment pupil dilation and potentially enhance sensorimotor processing by activating the locus coeruleus–norepinephrine (LC–NE) system. First, we characterized the temporal dynamics of the tVNS-induced pupillary response and developed a predictive model using a long short-term memory (LSTM) network in standard lighting. The model demonstrated efficacy in reproducing nonlinear physiological characteristics even under sudden dark exposure. Second, we derived optimized stimulation sequences using a genetic algorithm to maximize pupil dilation during immediate adaptation to darkness. Pupillary evaluations during bright-to-dark transitions revealed that the optimized tVNS additively increased pupil diameter. The improvement in visuomotor choice-reaction performance was highly correlated with the amount of tVNS-induced increase in pupil diameter during dark transition across healthy young adults. These findings suggest that the model-optimized tVNS approach may effectively mitigate the degradation of visuomotor performance during dark transitions via LC–NE activation as well as subsequent sensorimotor enhancement. Furthermore, the experimental results demonstrate the potential of the transient pupil response to tVNS to serve as a robust physiological indicator for predicting behavioral improvement.

## I. Introduction

Rapid changes in ambient illuminance are prevalent in operational environments, such as entering a tunnel or monitoring instruments during night flights. The human visual system undergoes dark adaptation to adjust to these low-light conditions [1]. During this adaptation process, visual information processing is temporarily impaired, which can lead to delayed behavioral responses and an increased risk of traffic accidents or operational errors [2], [3]. Dark adaptation in vision is mediated by two mechanisms: photoreceptor sensitivity and pupil dilation [4]. Since the pupil responds on a shorter time scale than the photoreceptors, its role is essential during early dark adaptation when behavioral performance is unstable. The importance of the pupillary response is particularly critical under conditions associated with degraded dark adaptation, such as aging, fatigue, and sleep deprivation [5]–[Maintaining or enhancing the speed of pupil dilation and associated visual information processing and motor execution is therefore essential for safety in environments with rapidly changing lighting.

A potential human–machine interface (HMI) approach to facilitate behavioral responses during dark adaptation is to artificially enhance locus coeruleus–norepinephrine (LC–NE) activity to induce optical and neurophysiological benefits. First, as an optical effect, pupil dilation increases retinal illuminance. Pupil diameter serves as a reliable biomarker for LC activity, as neurophysiological recordings have shown that LC firing consistently precedes pupil dilation [8], [9]. Second, regarding neurophysiological effects, LC–NE activity modulates neural sensory gain according to the adaptive gain theory [10], [11]. Specifically, LC–NE activation has been shown to enhance perceptual sensitivity across auditory [12], tactile [13], and visual [14] modalities. These physiological effects are linked to improved behavioral performance, including enhanced contrast detection and faster target identification [15], [16]. These findings suggest that LC–NE activation can concurrently improve visual cognitive performance through both increased retinal illuminance and enhanced neural processing. The LC–NE system can also be activated artificially by electrically stimulating the afferent vagus nerve due to their neuroanatomical connections. Stimulation of the cervical vagus nerve via an implanted electrode increases noradrenergic discharges, pupil diameter, and spontaneous motor activities including whisking and locomotion in rodents [17]. Collectively, artificial activation of the LC–NE system via vagus nerve stimulation may facilitate not only cognitive [18] but also motor performance.

As an alternative to the invasive approach, transcutaneous vagus nerve stimulation (tVNS) of the auricular branch of the afferent vagus nerve provides non-invasive access to the LC–NE system. Auricular stimulation activates brainstem pathways projecting to the LC [19]–[thereby modulating motor output gain [23] and sensory feedback integration [24]. Previous studies demonstrate that tVNS can enhance motor learning, facilitate action reprogramming, and shorten reaction times, particularly during action planning in challenging conditions [25]–[28]. Additionally, phasic tVNS evokes a rapid, transient pupil dilation, which is expected to temporarily increase retinal illuminance [29]. Under sudden light reductions, this transient pupil dilation may interact additively with the pupillary light reflex (PLR) [30], potentially mitigating performance degradation.

The objective of this study is to examine the efficacy of tVNS for visuomotor choice-reaction responses during bright-to-dark transitions in healthy young adults. As a first exploration into this fundamental function, we focus on developing and utilizing a tVNS-induced pupil response model and understanding the contribution of tVNS in a basic population and condition, without the confounding influences of aging, fatigue, or sleep deprivation. Specifically, we investigate whether tVNS can accelerate and augment pupillary dilation and influence choice-reaction performance in a visuomotor task performed at the onset of darkness. In addition, we assess whether tVNS-induced pupil dilation functions additively with the PLR under dynamically changing illumination, thereby validating response models developed under standard lighting conditions. Finally, we explore individual variability in visuo-motor responsiveness and task performance, with the aim of providing a foundation for personalized and adaptive HMIs.

## II. Experiment I: Relationships between tVNS and pupil response

Experiment I was conducted to model the pupil response to tVNS in a stable ambient environment and derive an optimal tVNS sequence to be used for testing its efficacy in dark adaptation in Experiment II.

### A. Participants and Experimental Setup

#### 1) Participants, Apparatus, and Environment

Twelve healthy, right-handed young adults participated (six males, six females; age: 22.6*±* 2.1 years, range 19–26). None reported fatigue or sleep deprivation. The protocol was approved by the Institutional Review Board at the Georgia Institute of Technology (Protocol: IRB2025-104), and written informed consent was obtained from all participants.

Monophasic rectangular pulses (30 Hz, pulse width: 100 *µ*s, duration time: 0.5 s; Fig. 1(a)) were generated by an isolated constant-current stimulator (DS7AH, Digitimer Ltd., UK) controlled via a microcontroller (Raspberry Pi Pico). As shown in Fig. 1(b), active tVNS was delivered to the anterior and posterior left tragus using hydrogel electrodes (RELIfit-Tragus, Soterix Medical, NJ, USA), while Sham stimulation was applied to the left earlobe (Easy-Clip, Soterix Medical). Pupil diameter was recorded binocularly at 60 Hz using a screen-based eye tracker (Tobii Pro Spark, Tobii Corp., Sweden). Visual stimuli programmed in PsychoPy [31] were presented on a calibrated 27-inch LCD monitor (54 cd/m^2^, DELL P2722H). Participants sat with their head stabilized by a chin rest at a 56-cm viewing distance under 500-lux ambient illuminance, fixating on a central red circle (diameter: 35 pixels, visual angle: *∼*1.1^*°*^).

**Fig. 1.**
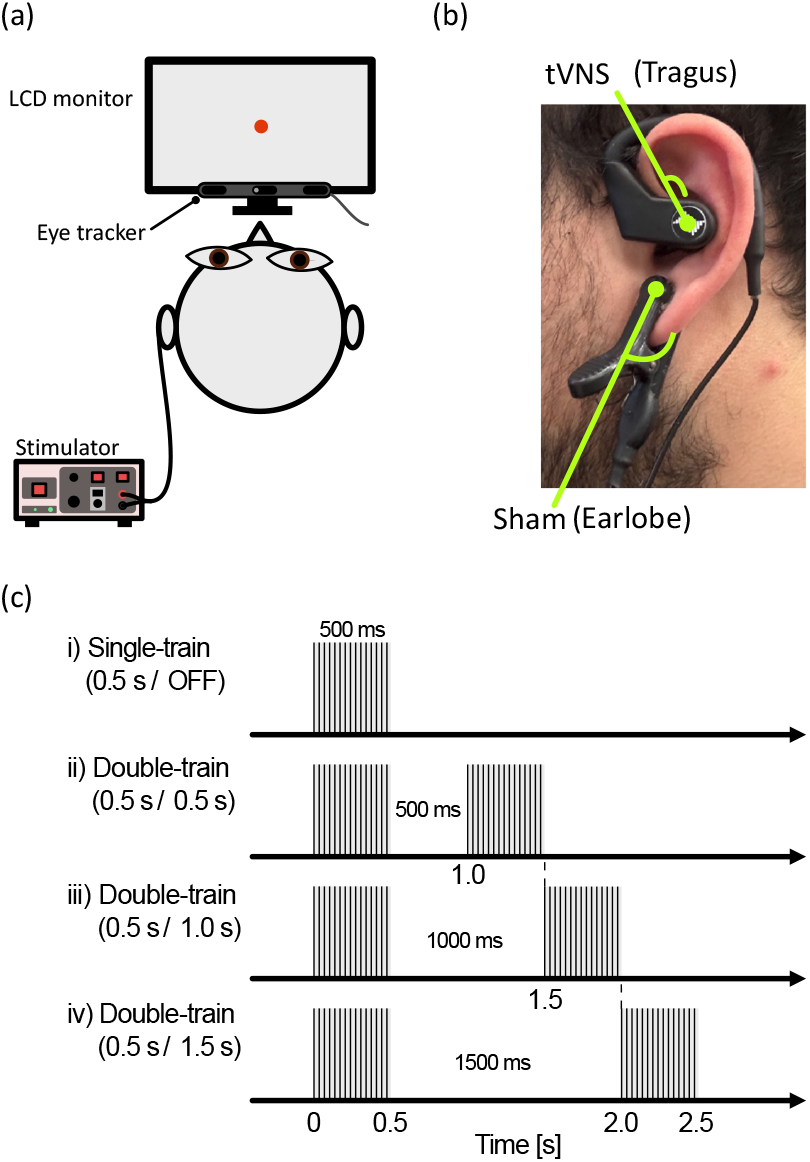
Experimental setup for Experiment I: (a) schematic representation of the system, where electrical pulses are delivered via the stimulator while pupillary responses are recorded, and (b) electrode placement for the transcutaneous stimulation. The active electrode for tVNS is positioned on the tragus, while the Sham electrode is attached to the earlobe. (c) stimulation sequences. 30 Hz pulsed stimulations are delivered within a 0.5 s window. The time interval between the first and second stimuli is varied.

#### 2) Stimulation Protocol and Calibration

Current intensity was calibrated individually, capped strictly at 5.0 mA. Active tVNS was set 0.2 mA below the perceived pain threshold. Sham intensity was matched to tVNS if it elicited a stable cutaneous sensation, or adjusted to the minimum level providing stable sensation below the Sham pain threshold. Four stimulation sequences were tested (Fig. 1(c)): single-train (0.5 s / OFF) and double-trains with inter-train intervals of 0.5, 1.0, or 1.5 s (*T*_ON_ = 0.5 s). Each participant completed 12 alternating blocks of tVNS and Sham (48 trials total, 12 per sequence in pseudorandom counterbalanced order). Each 10-s trial (3.0 s pre-stimulation, 7.0 s post-stimulus) was separated by at least 90 s of rest for pupillary recovery.

#### 3) Signal Preprocessing and Feature Extraction

Pupil data were cleaned using a custom algorithm detecting blinks, eyetracker validity flags, and physiological anomalies (*>* 0.2 mm frame-to-frame changes). Gaps *<* 2.0 s were linearly interpolated. Only two Sham trials in one participant were excluded due to continuous signal loss, yielding an overall valid trial rate of 99.65% (574/576 trials; tVNS: 100.0%, Sham: 99.31%). Data were baseline-normalized to compute the pupil dilation response (PDR) as *z*(*t*) = 100 *×* [*x*(*t*) *−µ*_base_]*/µ*_base_ [%], where *µ*_base_ denotes the 1.0-s prestimulus baseline mean. Within a 3.0-s post-stimulus analysis window, three physiological features were extracted: Area under the Curve (AuC, time integral from 0 to 3 s), peak amplitude (maximum percentage change), and duration continuously exceeding 50% of the peak amplitude (*D*_50%_).

#### 4) Statistical Analysis

Statistical analyses were performed using MATLAB (MathWorks, Inc., MA, USA) equipped with the Statistics and Machine Learning Toolbox and the FieldTrip toolbox [32]. The significance level was set at *α* = 0.05 for all tests.

##### Time-Series Analysis

To evaluate the time-course differences between tVNS and Sham conditions without making a priori assumptions about the timing of the effect, we employed a cluster-based permutation test. This non-parametric method controls for the multiple comparison problem inherent in time-series data by clustering adjacent time points that exhibit significant differences (*p <* 0.05) and comparing the sum of their t-statistics against a null distribution generated from 1,000 random permutations.

##### Scalar Feature Analysis (Response Magnitude)

Pupil response characteristics (AuC, peak amplitude, and *D*_50%_) were analyzed using a linear mixed-effects model (LMM) incorporating stimulation type (tVNS vs. Sham), stimulation sequence (four sequences), and their interaction as fixed effects, with participant as a random intercept. Holm–Bonferroni correction was applied for multiple comparisons across the response variables.

### B. Results

#### 1) Time-series analysis

To determine the temporal extent of the tVNS-specific effect, we compared the pupil dilation response profiles between tVNS and Sham stimulations using a cluster-based permutation test. Fig. 2(a)–(d) illustrates the time-series comparison for each stimulation sequence. Significant clusters appeared in all but the double-train (0.5 s / 1.0 s) sequence. For example, in the double-train (0.5 s / 0.5 s) sequence, a significant positive cluster (*p <* 0.042) was identified extending from *t* = 1.6 s to *t* = 2.6 s after the onset of stimulation, indicating that tVNS elicited a significantly greater pupil dilation than Sham during this time window. Similar clusters were observed in the other sequences, confirming that tVNS reliably elicited pupillary dilation across different stimulation sequences.

**Fig. 2.**
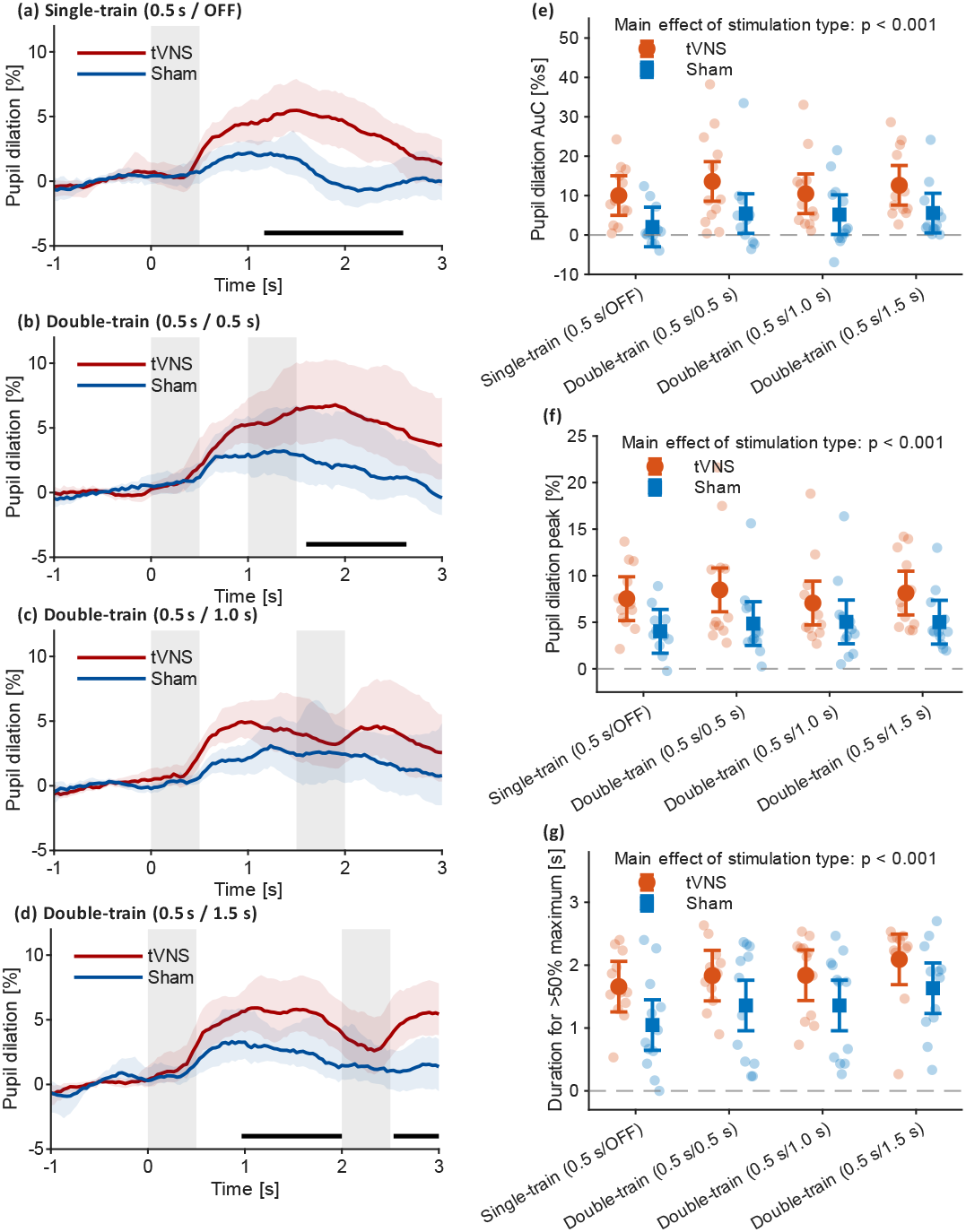
Pupil diameter responses and response magnitude characteristics for tVNS and Sham across varying stimulation sequences in Experiment I. (a)–(d) Time courses of baseline-normalized pupil dilation responses for (a) single-train (0.5 s / OFF), (b) double-train (0.5 s / 0.5 s), (c) double-train (0.5 s / 1.0 s), and (d) double-train (0.5 s / 1.5 s) sequences. Red and blue lines represent grand-mean responses for tVNS and Sham, respectively; shaded regions denote 95% confidence intervals (CIs). Gray rectangles indicate the timing and duration of stimulation trains (*T*_ON_ = 0.5 s). Horizontal black bars indicate time intervals with significant differences between tVNS and Sham (*p <* 0.05, cluster-based permutation test). (e)–(g) Physiological response metrics across stimulation sequences: (e) Area under the Curve (AuC), (f) peak dilation amplitude, and (g) duration continuously exceeding 50% of the peak amplitude (*D*_50%_). Light red and blue dots represent individual-participant data for tVNS and Sham, respectively. Solid symbols and error bars represent linear mixed-effects model (LMM) estimated means and 95% CIs. A significant main effect of stimulation type was observed for all three measures (*p <* 0.001).

#### 2) Pupil response magnitude

To represent the overall and peak magnitude of pupil dilation across the response period, AuC and peak amplitude over the entire 3 s served as the dependent variables, respectively. A significant main effect of stimulation type was found for AuC (*F* (1, 88) = 35.19, *p <* 0.001, *d* = 1.21), peak amplitude (*F* (1, 88) = 34.26, *p <* 0.001, *d* = 1.19), and *D*_50%_ (*F* (1, 88) = 17.46, *p <* 0.001, *d* = 0.85), confirming that tVNS induced consistently greater pupil dilation than Sham stimulation across all sequences as shown in Fig. 2(e)–(g). After Holm–Bonferroni correction, no significant main effect of stimulation sequence was observed for any measure (AuC: *F* (3, 88) = 1.70, *p* = 0.173; peak amplitude: *F* (3, 88) = 0.65, *p* = 0.586; *D*_50%_: *F* (3, 88) = 2.95, *p* = 0.037). No significant interaction between stimulation type and sequence was found for any measure (AuC: *F* (3, 88) = 0.30, *p* = 0.827; peak amplitude: *F* (3, 88) = 0.48, *p* = 0.698; *D*_50%_: *F* (3, 88) = 0.08, *p* = 0.968).

## III. Stimulation sequence optimization

### A. Problem formulation

The pupil control system regulated by tVNS is modeled as a single-input single-output (SISO) discrete-time dynamic system. We define the electrical stimulation parameters as the input *u*(*k*) and the observed pupil dilation response as the output *z*(*k*), where *k* denotes the sample index. To focus on the relatively slow temporal dynamics of the pupillary response and reduce computational cost, the preprocessed data were downsampled to 10 Hz.

Recent research has shown that both phasic and tonic tVNS evoke rapid but transient pupil dilation, suggesting that the temporal characteristics of these responses are relatively consistent across stimulation durations [28]. Our models aim to capture these nonlinear dynamics. The models were fitted using data from *−*2.9 s to +7.0 s relative to stimulus onset, and predictive performance was evaluated from *−*1 s to +3 s relative to stimulus onset. We evaluated and compared two representative modeling approaches: an autoregressive with exogenous inputs (ARX) model and a nonlinear long short-term memory (LSTM) network.

### B. Models for pupil responses

#### 1) Linear model (ARX)

As a linear approach, we constructed an ARX model described by the following difference equation:

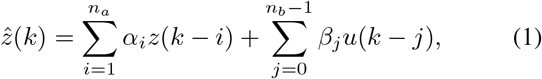

where 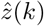 is the estimated pupil response, *z*(*k*) is the observed response, and *u*(*k*) is the exogenous input, and *α*_*i*_ and *β*_*j*_ are the model coefficients to be estimated. The optimal model orders *n*_*a*_ and *n*_*b*_ were determined by minimizing the Akaike information criterion (AIC).

#### 2) Nonlinear model (LSTM)

To capture nonlinearities and long-term temporal dependencies, we utilized an LSTM network. Our model was implemented using a stacked LSTM architecture consisting of two LSTM layers, each with 100 hidden units. To prevent overfitting, a dropout layer with a rate of 0.02 was placed before the final dense output layer. This final layer employed a linear activation function. The loss function was defined as the Mean Squared Error (MSE) between the predicted and observed pupil response, supplemented by *L*_2_ regularization.

#### 3) Input data filtering

The tVNS stimulation was applied in the form of a pulse train (pulse width: 100 *µ*s, frequency: 30 Hz, duration time: 0.5 s). Considering the sampling rate of 10 Hz and the fact that biological tissue response time constants are much longer than individual electrical pulses, we defined the discrete input *u*(*k*) using two distinct representations:

- *Binary envelope:* Defining *u*(*k*) as the binary envelope (On/Off state) of the pulse train. To synchronize the input signal with this 10-Hz temporal grid, the high-frequency pulse train (30 Hz) was mapped onto discrete steps by assigning *u*(*k*) = 1 if the corresponding 100-ms interval contained stimulation pulses, and 0 otherwise.
- *Biomimetic filter:* Neural responses are characterized by specific physiological properties: smoothing (integration of short-term stimuli), adaptation (diminishing response to sustained stimuli), and rectification (non-negative firing rates). Following mathematical models of the PLR [33], we defined a filtered input signal *u*_phys_(*k*) by combining linear filters and static nonlinear elements. The transient response *u*_AC_(*k*), representing adaptation, was modeled using a high-pass filter defined by the transfer function 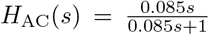, which represents how the system responds to rapid changes while filtering out steady-state components. Finally, half-wave rectification was applied as *u*_phys_(*k*) = max {0, *u*_AC_(*k*)} to reflect the non-negativity of neural firing rates.

### C. Model performance evaluation

For model identification and evaluation, only the tVNS trials were used. Of the 576 trials recorded in Experiment I, 288 were tVNS trials and 288 were Sham trials. For each participant, the six repeated tVNS trials for each of the four stimulation sequences were averaged after preprocessing, yielding 48 participant-by-sequence mean waveforms (12 participants *×* 4 sequences) for modeling. Thus, each model input–output observation was an averaged waveform rather than a single raw trial.

To evaluate the predictive performance and generalization capability of the models to unseen stimulation waveforms, we used leave-one-stimulation-sequence-out cross-validation. For each fold, 36 participant-by-sequence mean waveforms from three stimulation sequences (12 participants per sequence) were used to fit the models, while the 12 participant-by-sequence mean waveforms from the remaining sequence were used for evaluation, ensuring that the held-out stimulation sequence was unused during training.

For the ARX model, the model orders were optimized by searching for the autoregressive order *n*_*a*_ and exogenous input order *n*_*b*_ within the range of 1 to 15 to minimize the AIC calculated on the training dataset. The selected models had *n*_*a*_ values ranging from 3 to 11 and *n*_*b*_ values ranging from 6 to 10. For the LSTM models, each configuration was trained using five predefined random seeds per fold and input type, with predictive performance calculated for each run and then averaged across runs.

The predictive performance of the models was evaluated over the temporal window from *−*1 s to +3 s relative to stimulus onset, using the root mean square error (RMSE), mean absolute error (MAE), and coefficient of determination (*R*^2^). Table I summarizes the performance across four configurations: ARX versus LSTM models, and binary input (*u*) versus biomimetic input (*u*_phys_).

**Table I.** Comparison of Model Performance.

| Model | Input type | RMSE | MAE | $R^2$ |
| --- | --- | --- | --- | --- |
| ARX | Binary ($u$) | $1.20 \pm 0.18$ | $0.91 \pm 0.12$ | $0.68 \pm 0.15$ |
| ARX | Biomimetic ($u_{\text{phys}}$) | $1.21 \pm 0.18$ | $0.91 \pm 0.14$ | $0.68 \pm 0.12$ |
| LSTM | Binary ($u$) | $0.95 \pm 0.32$ | $0.78 \pm 0.22$ | $0.80 \pm 0.10$ |
| LSTM | Biomimetic ($u_{\text{phys}}$) | $0.98 \pm 0.37$ | $0.82 \pm 0.28$ | $0.79 \pm 0.11$ |

As reported in Table I, the LSTM models showed better predictive performance than the ARX models for both input types. The LSTM model using the biomimetic input achieved comparable predictive performance with binary input. These results support the ability of the LSTM models to predict participant-averaged pupillary response waveforms for previously unseen stimulation conditions.

### D. Stimulation Sequence Optimization

Due to constraints related to the project timeline, Experiment II had to be completed within a predetermined timeframe. To accommodate this requirement, Experiment II commenced immediately after the model development from the dataset of the initial five participants. In this process, the model derived from the five participants was applied to compare the predicted pupil responses with measured ones, derive an optimal sequence for dark transition, and compare the pupil response predicted from the derived sequence with those from the employed sequences. After Experiment II was completed, we compared the predicted AuC between the models developed using data from 5 and 12 participants.

#### 1) Formulation of Objective Function

To visualize predicted profiles, Fig. 3(a) overlays the predictions generated by the LSTM model using the biomimetically filtered input (*u*_phys_), fitted to those participants, with the measured pupil responses averaged across all twelve participants. The demonstration includes the four tested stimulation sequences: Single (0.5 s / OFF), Double (0.5 s / 0.5 s), Double (0.5 s / 1.0 s), and Double (0.5 s / 1.5 s) trains. This model was used to derive the stimulation sequence in Experiment II.

**Fig. 3.**
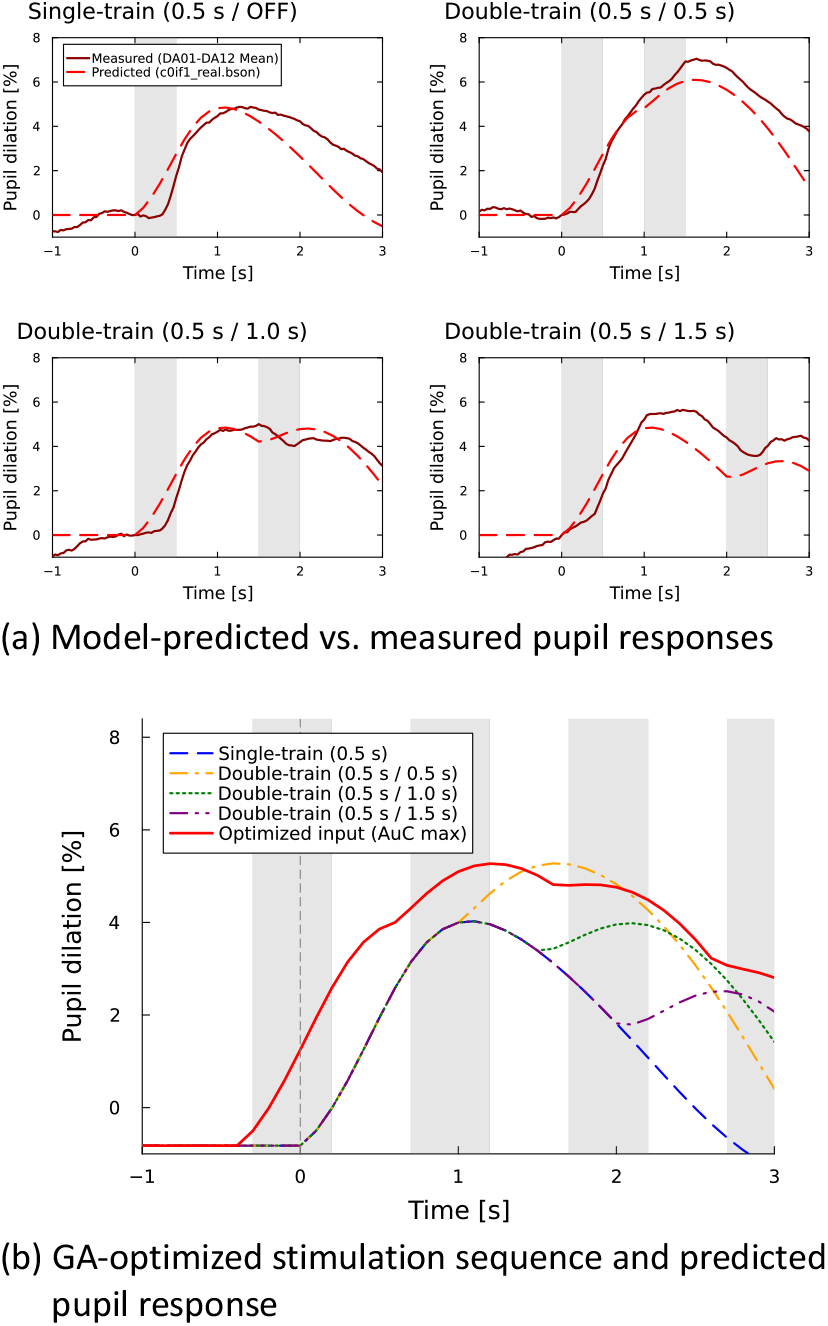
Model validation and stimulation sequence optimization. (a) Comparison between model-predicted (dashed red lines) and measured pupil dilation responses (brown lines; mean across all twelve participants) across four tested stimulation sequences. (b) Estimated pupil response to the GA-optimized tVNS sequence (solid red line) compared with the tested sequences (dashed/dotted lines).

The model predicted the discrete pupil response 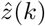 for each candidate stimulation sequence at 10 Hz. The onset of darkness is defined as *k* = 0. The objective function *J* is designed to minimize the pupil diameter before the onset of darkness while maximizing it afterward to facilitate dark adaptation. We employ an unscaled discrete objective proportional to the difference in the AuC of the pupil diameter before and after the onset of darkness as follows:

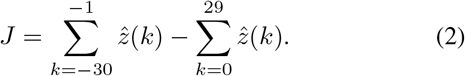

A window of *±*3.0 s (30 samples on each side) was selected to encompass the typical duration of the target detection task and to cover the critical phase of the dark adaptation process. The common sampling interval was omitted from *J* because it only scales all objective values by the same constant and therefore does not affect the minimizing sequence. In this formulation, the first term represents the cumulative predicted response during the 3 s preceding the onset of darkness, which serves as a penalty because pupil dilation before an illuminance change can prolong the subsequent dark adaptation period. The optimization task aims to find an input vector *u* that minimizes *J* (thereby maximizing the relative dilation after *k* = 0).

#### 2) Derivation of Optimal Sequence via Genetic Algorithm

We obtained the optimized input vector *u*_opt_ using a genetic algorithm (GA). Each element of the input vector *u* is binary (*u*_*n*_ ∈{0, 1}), and the vector was used directly as the chromosome. Based on the preliminary experimental results, the minimum continuous OFF duration and continuous ON duration were both set to 0.5 s as a restriction condition. Consequently, the chromosome contained 60 samples (6 s at 10 Hz).

The GA was configured with a population size of 100, 500 generations, a crossover rate of 0.7, and a mutation rate of 0.05. Specifically, any sequence of consecutive ON (ones) or OFF (zeros) states shorter than 0.5 s (5 consecutive bits) was prohibited. The optimization process was executed to minimize the objective function *J* in (2). During the evolution, individuals that failed to meet the temporal constraints were either assigned a high penalty value to their fitness score or were adjusted to the nearest valid stimulation sequence.

The optimization results are presented in Fig. 3(b). The optimal tVNS sequence began 0.4 s before the onset of darkness (*t* = 0), providing feedforward compensation for the physiological latency observed in empirical pupil responses. The optimal Sham sequence began 0.6 s before darkness, consistent with its slower response dynamics. The estimated AuC produced by the optimized tVNS sequence was 12.32 %*·*s, exceeding those produced by the single- and double-train sequences (5.11, 9.64, 8.70, and 7.26 %*·* s). The optimized Sham sequence produced an estimated AuC of 7.08 % *·*s, exceeding the corresponding values for the tested sequences (2.72, 5.15, 4.46, and 3.71 %*·* s).

In Table 1 analysis, the performance of LSTM-Biomimetic and Binary models was similarly high. However, the optimized stimulation sequence was different between two filters. Optimization for the LSTM-Binary model outputted a single and long pulse, whereas the LSTM-Biomimetic model outputted multiple short pulses at set intervals. Previous studies have shown that pupil dilation can be elicited by stimulation trains as short as approximately 0.6 s, while extending the stimulation duration does not necessarily sustain the response [28], [34]. This supports our selection of the Biomimetic model for stimulation sequence optimization in Experiment II. With Biomimetic input, there was only a small difference in AuC between the models developed using data from 5 and 12 (12.58 % *·*s) participants, suggesting that the employed stimulation sequence was acceptable

## IV. Experiment II: Efficacies of tVNS during dark adaptation

Experiment II examined whether the model-optimized tVNS sequence enhanced visuomotor performance during the transition from bright to dark conditions, as assessed by pupillary responses and manual choice-reaction performance. Because data collection for Experiments I and II partially overlapped, the sequence was derived from the Experiment I data available from the first five participants. Once determined, the sequence was fixed and applied without modification to all twelve participants in Experiment II.

### A. Participants and Experimental setup

#### 1) Participants and Environment

The same 12 healthy young adults from Experiment I participated in Experiment II, and the identical electrical stimulation and eye-tracking systems were employed. The experiment was conducted in a controlled twilight environment with ambient illuminance maintained at 50 lux around the participants’ eyes.

#### 2) Task Protocol and Experimental Design

The visuo-motor choice-reaction task required participants to respond to a circular target (diameter: 1.9^*°*^, 60 pixels; luminance: 0.4 cd/m^2^) presented at one of eight randomized radial positions at an eccentricity of 8.0^*°*^ (250 pixels, 45^*°*^ intervals). As illustrated in Fig. 4(a), each trial comprised three consecutive phases: (1) *Fixation (7 s):* Central red fixation point (35 pixels) on a gray background (54 cd/m^2^). (2) *Bleaching (3 s):* White background (250 cd/m^2^) to induce temporary retinal bleaching, with the fixation point set to light gray. (3) *Task (7 s):* Black background (0.27 cd/m^2^) with target presentation, where participants moved the cursor from the center to click the target as rapidly as possible using a mouse with their right hand.

**Fig. 4.**
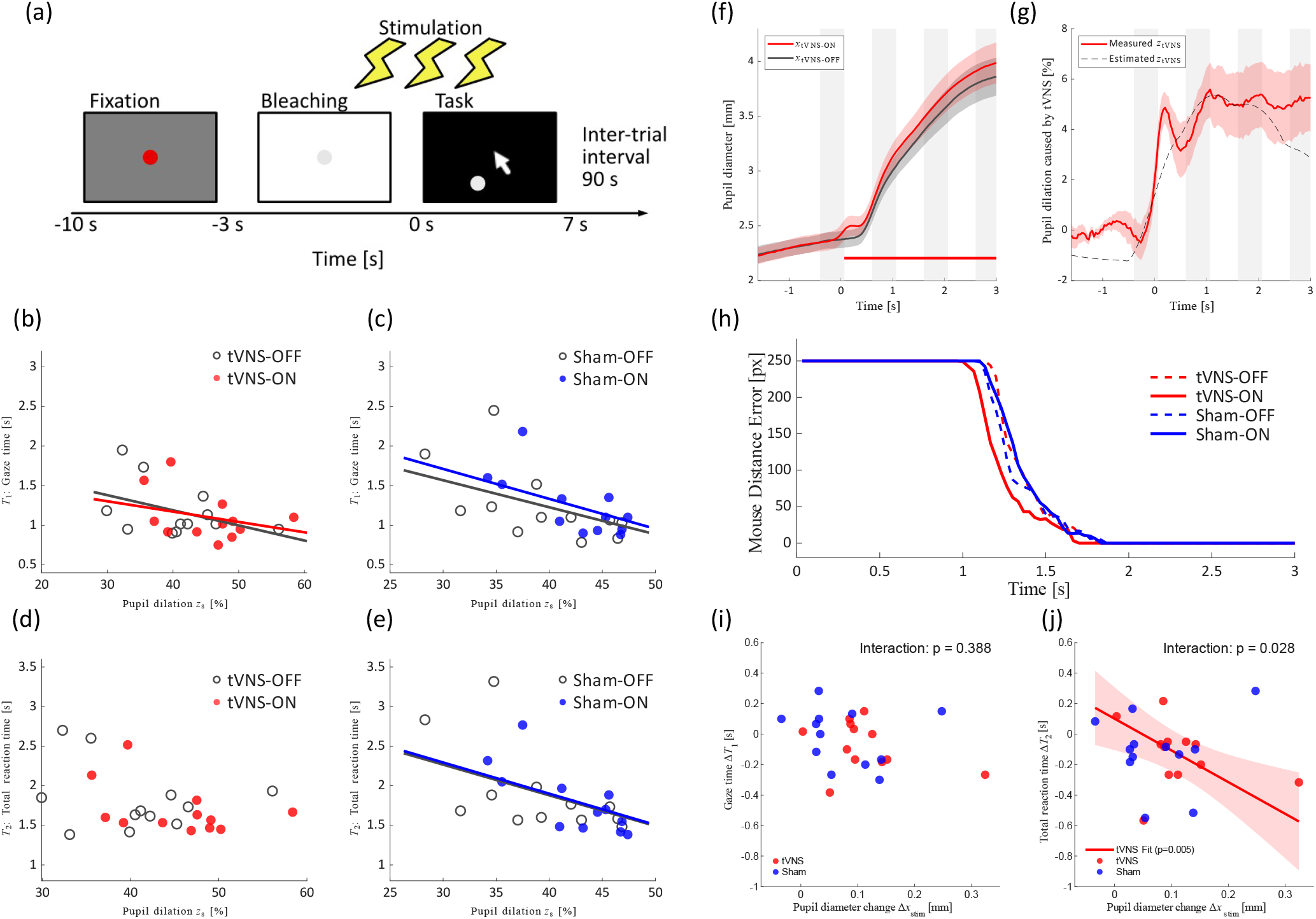
Experimental protocol, pupillary dynamics, and visuomotor choice-reaction performance during dark transition in Experiment II. (a) Timed sequence of a single trial comprising Fixation, Bleaching, and Task phases, with model-optimized stimulation delivered just prior to dark transition. (b)–(e) Visuomotor performance as a function of grand-mean centered pupil dilation: (b) gaze time (*T*_1_) in tVNS, (c) *T*_1_ in Sham, (d) mouse-click total reaction time (*T*_2_) in tVNS, and (e) *T*_2_ in Sham. Filled circles represent stimulation-ON (red: tVNS; blue: Sham), open circles denote stimulation-OFF, and solid lines indicate significant linear mixed-effects model (LMM) fits. (f) Grand-average time courses of absolute pupil diameter during dark adaptation with and without tVNS (the horizontal bar denotes cluster *p <* 0.05). (g) Additive property demonstrated by comparing the empirical tVNS-induced pupil component during dark transition (solid red line) with the model-predicted response under static lighting (dashed black line). (h) Temporal profiles of mouse distance error relative to the target across stimulation states. (i), (j) Relationships between stimulation-induced pupil dilation (Δ*x*_stim_) and performance changes: (i) change in gaze time (Δ*T*_1_) and (j) change in total reaction time (Δ*T*_2_). Red and blue circles represent participant-level data for tVNS and Sham, respectively; shaded bands indicate 95% confidence intervals of LMM fits.

The model-optimized stimulation (*u*_opt_) was delivered prior to the dark transition according to the lead time identified in Section III (0.4 s for tVNS and 0.6 s for Sham), with current intensities calibrated identically to Experiment I. The session comprised two 16-trials (32 trials total). One session included tVNS-ON and tVNS-OFF, and the other included Sham-ON and Sham-OFF. Each session contained eight ON and eight OFF state trials, with each of the eight target positions presented twice in pseudorandom order. Session order was counterbalanced across participants. An inter-trial interval of at least 90 s was strictly maintained for physiological recovery and minimizing fatigue.

#### 3) Performance Variables

To evaluate the effect of tVNS on sensorimotor choice-reaction performance, we defined two temporal variables relative to the onset of target appearance (*t* = 0):

- *Gaze time (T*_1_): This metric represents the latency of the initial saccadic response toward the presented target based on the eye-tracker data. To ensure robust landing detection against ocular noise or blinks, we implemented a backward-tracking algorithm. First, gaze velocity was calculated by the discrete derivative of the Euclidean distance between the gaze position and the target center. Subsequently, the algorithm identified the last point where the velocity exceeded a predefined threshold (15^*°*^*/*s) within the response window and then searched backward to find the earliest continuous point of threshold crossing, defining it as gaze landing. The time from target onset to the detected gaze landing was defined as gaze time.
- *Total reaction time (T*_2_): This metric reflects the total task completion time, measured from mouse position recording. The task completion was defined as the moment when a successful mouse clicking (logical high) was recorded while the cursor was within the target.

We employed the median value across trials for these temporal variables (*T*_1_ and *T*_2_) to represent the performance of each participant.

#### 4) Pupil Response Metrics

Pupil data were preprocessed and noise-reduced in a similar manner to Experiment I. One trial (tVNS-OFF) was excluded from the analysis in one participant because of continuous data loss exceeding 2.0 s (resulting in less than 50% valid pupil samples for that trial), yielding an overall valid trial rate of 99.74% (383 of 384 trials across all participants). Following artifact removal, the data were trigger-averaged to the task onset across the valid trials for each stimulation type and state combination.

The pre-task diameter before the onset of each trial was established as the mean pupil diameter from *−*1.6 s to *−*0.6 s prior to the onset of darkness (*t* = 0): 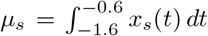 The mean pupil dilation during the task relative to this pretask diameter, *z*_*s*_, was defined in relative terms [%] under state *s ∈ {*tVNS-ON, tVNS-OFF, Sham-ON, Sham-OFF*}*, averaged over the initial 3.0 s window as 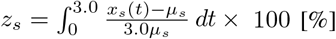, where *x*_*s*_(*t*) [mm] is the absolute pupil diameter in state *s*. The integrals were approximated discretely at 30 Hz (Δ*t ≈* 0.033 s), and *z*_*s*_ is expressed as a percentage.

To quantify the partial enlargement of pupil diameter artificially induced by the tVNS or Sham stimulation specifically, we defined the part of pupil diameter change caused by stimulation Δ*x*_stim_. This metric represents the difference between the mean pupil responses across 0–3 s between stimulation-ON and -OFF states in mm. Let *x*_stim-ON_(*t*) and *x*_stim-OFF_ (*t*) denote pupil diameter under the stimulation-ON and stimulation-OFF states, respectively. By subtracting the pupil diameter under the stimulation-OFF state from that under the stimulation-ON state, the metric is defined as 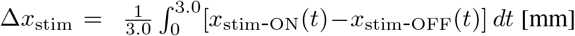. A positive value in this metric indicates that the delivery of model-optimized stimulation induced greater pupil dilation compared to the non-stimulated state. Specifically, we denote this metric as Δ*x*_tVNS_ when evaluating the effects of active tVNS (using *x*_tVNS-ON_ (*t*) and *x*_tVNS-OFF_ (*t*)) and Δ*x*_Sham_ when evaluating those of Sham (using *x*_Sham-ON_ (*t*) and *x*_Sham-OFF_ (*t*)).

The changes in choice-reaction performance were calculated as the differences in median values between the stimulation and no-stimulation states for each participant. Specifically, the change in gaze time (Δ*T*_1_) was defined as Δ*T*_1_ = *T*_1,stim-ON_ *−T*_1,stim-OFF_, and the change in total reaction time (Δ*T*_2_) was defined as Δ*T*_2_ = *T*_2,stim-ON_ *−T*_2,stim-OFF_.

The magnitude of the natural PLR, independent of electrical stimulation, was defined as *z*_PLR_ and calculated as *z*_*s*_ in the Sham-OFF condition. A greater *z*_PLR_ [%] indicates greater mean pupil dilation during the initial phase following a sudden reduction in illumination. The Sham-OFF condition was used to quantify natural pupillary responsiveness because no electrical stimulation was delivered during these trials and no active tVNS was administered within the Sham session, thereby minimizing potential carryover effects of vagal stimulation.

#### 5) Statistical Analysis

To investigate associations among stimulation state, mean pupil dilation, and visuomotor choice-reaction performance within each stimulation type, we used linear mixed-effects models (LMMs). Separate models were fitted for the tVNS and Sham sessions, with either gaze time (*T*_1_) or total reaction time (*T*_2_) as the dependent variable. Fixed effects included stimulation state (ON vs. OFF), mean pupil dilation, and their interaction. A participant-specific random intercept accounted for repeated observations and inter-individual differences. Mean pupil dilation was grandmean centered separately within each session before model fitting so that the intercept and main effect of stimulation were evaluated at the mean pupil-dilation level.

Mediation analyses were also conducted separately for the tVNS and Sham sessions and for gaze time (*T*_1_) and total reaction time (*T*_2_). Stimulation state was coded as OFF = 0 and ON = 1. For each participant and condition, reaction times were summarized as the median across trials. Mean pupil dilation was calculated from baseline-normalized trial responses by averaging over the predefined post-stimulation window and across valid trials and was grand-mean centered before analysis. Three linear mixed-effects models with a participant-specific random intercept estimated path *a* (pupil dilation regressed on stimulation state), paths *b* and *c*^*′*^ (reaction time regressed on pupil dilation and stimulation state), and the total effect *c* (reaction time regressed on stimulation state). No stimulation-by-pupil-dilation interaction was included, allowing the standard decomposition *c* = *c*^*′*^ +*ab*. The indirect effect *ab* was evaluated using the first-order delta (Sobel) method, with two-sided *p*-values and 95% confidence intervals.

To compare the temporal profile of the pupil diameter response as well as cursor trajectories between conditions, cluster-based permutation tests were applied. The procedure included an initial threshold for cluster identification and a final threshold for statistical comparison.

The efficacy of tVNS for visuomotor performance was investigated by examining the relationship between the change in visuomotor performance variables and the change in pupil diameter specific to stimulation. To compare such relationships between stimulation types (tVNS vs. Sham), we used LMMs. In these models, the changes in performance (gaze time Δ*T*_1_ and total reaction time Δ*T*_2_) were the dependent variables. The fixed effects included the stimulation type (tVNS vs. Sham), the pupil dilation caused by stimulation (Δ*x*_stim_), and their interaction (representing the difference in slopes between stimulation types). Δ*x*_stim_ (Δ*x*_tVNS_ and Δ*x*_Sham_) was group-mean centered within each stimulation type prior to model fitting. A random intercept for each participant was also included. Slopes and their 95% CIs for each condition were estimated using the fixed-effects covariance matrix and the residual degrees of freedom.

To further assess whether the changes in visuomotor choice-reaction time due to electrical stimulation differed from zero (i.e., performance change), Δ*T*_1_ and Δ*T*_2_ were evaluated using one-sample *t*-tests for each stimulation type. Additionally, analysis with a multivariate LMM was performed, in which the change in total reaction time (Δ*T*_2_) was set as the dependent variable. Stimulation type (tVNS vs. Sham, with Sham as the reference), stimulation-induced pupil dilation response (Δ*x*_stim_), natural PLR (*z*_PLR_), interactions between stimulation type and Δ*x*_stim_, and interactions between stimulation type and *z*_PLR_ were included as fixed effects. A random intercept was applied for each participant. The significance level was set at *α* = 0.05 for all tests.

### B. Results

To inspect the motor response dynamics, cursor distance relative to the target was tracked over time (Fig. 4(h)). Reaching movements initiated earlier with tVNS (*∼*1.0 s) than with Sham or no-stimulation conditions (*∼*1.15 s), exhibiting a transient cluster between 1.13 and 1.30 s (*p* = 0.116).

#### 1) Mean pupil dilation and reaction times

Pupil diameter was altered and variable during the task across participants, which may involve the modulating factors of PLR and electrical stimulation. Visuomotor choice-reaction performance was also variable across participants, ranging from 0.75–2.45 s for gaze time and from 1.38–3.32 s for total reaction time across stimulation types and states. The potential association across stimulation state, mean pupil dilation, and visuomotor performance among participants was investigated using the LMM analysis followed by mediation analysis for each reaction time within each stimulation type.

First, we examined the association of mean pupil dilation with stimulation state and stimulation type. The analysis found that mean pupil dilation was greater when the stimulation state was ON compared with OFF in both the tVNS session (tVNS-ON vs. tVNS-OFF; *b* = 4.74, *p <* 0.001) and the Sham session (Sham-ON vs. Sham-OFF; *b* = 3.38, *p <* 0.001).

We then examined whether mean pupil dilation across stimulation states was associated with visuomotor reaction performance. As shown in Fig. 4(b) and (c), mean pupil dilation was negatively associated with gaze time (*T*_1_) in both the tVNS session (*b* =*−* 0.019, *p* = 0.041) and the Sham session (*b* = *−*0.034, *p* = 0.017). For total reaction time (*T*_2_), as shown in Fig. 4(d) and (e), no significant association with mean pupil dilation was found in the tVNS session (*b* = *−*0.015, *p* = 0.197), whereas a significant negative association was found in the Sham session (*b* = *−*0.039, *p* = 0.028).

To examine whether mean pupil dilation mediated the association between stimulation state and *T*_1_, we performed a mediation analysis based on the LMM. The analysis yielded negative estimated indirect effects of stimulation state on *T*_1_ through mean pupil dilation in both sessions, although the effect reached nominal statistical significance only in the Sham session. In the Sham session, the pathway linking Sham-ON to *T*_1_ through mean pupil dilation yielded a significant negative indirect effect (*a × b* = *−*0.115 s, *Z* = *−*2.18, *p* = 0.030), despite a total-effect estimate close to zero (*c* = *−*0.018 s, *p* = 0.73). The corresponding direct-effect estimate was positive but not statistically significant (*c*^*′*^ = 0.097 s, *p* = 0.177). Thus, the opposing directions of the indirect and direct-effect estimates were consistent with a suppression pattern, although the evidence did not establish a positive direct effect. In the tVNS session, the indirect effect did not reach statistical significance (*a×b* =*−* 0.083 s, *Z* = *−*1.86, *p* = 0.064), but its estimated magnitude was similar to that of the total stimulation effect (*c* = −0.075 s, *p* = 0.108); the direct-effect estimate was close to zero (*c*^*′*^ = *−*0.008 s, *p* = 0.891).

After accounting for mean pupil dilation, stimulation state was not significantly associated with *T*_2_ in either session (tVNS: *c*^*′*^ = *−*0.057 s, *p* = 0.460; Sham: *c*^*′*^ = 0.018 s, *p* = 0.845). The LMM-based mediation analysis indicated that the indirect pathway through mean pupil dilation was significant in the Sham session (indirect effect: *a × b* = *−*0.131 s; *Z* = *−*2.05, *p* = 0.041; total effect: *c* = *−*0.113 s, *p* = 0.105) but not in the tVNS session (indirect effect: *a × b* = *−*0.076 s; *Z* = *−*1.40, *p* = 0.161), whereas the total effect of stimulation on *T*_2_ was significant (*c* = *−*0.133 s, *p* = 0.029).

#### 2) Pupil Responses Attributable to the Pupillary Light Reflex and tVNS

To examine additivity between the PLR and tVNS-induced dilation, we determined the contribution of tVNS in the time-course response of the absolute pupil diameter (Fig. 4(f)). Using the optimized sequence derived from the modeling with AuC analysis, the permutation analysis found a significant upward shift in pupil diameter with tVNS compared to the response without stimulation during the dark-adaptation period (*p* = 0.002). Subtracting the response without stimulation from that during tVNS at each time point yielded the instantaneous pupil response independent of the contribution of PLR. The additivity of two dilation mechanisms was then evaluated by comparing thus-derived pupil diameter change against the predicted response based on the model under nonvarying lighting (Fig. 4(g)). The calculated tVNS-induced pupil response independent of PLR showed a temporal profile comparable to the model-predicted tVNS-induced response, with a significant correlation (*r* = 0.92, *p <* 0.001). This observation supports additivity and suggests that tVNS effectively augmented pupillary dynamics and associated autonomic modulation even during the dynamic transient phase of dark adaptation.

To determine whether the tVNS-induced pupil response depended on participants’ natural light-reflex characteristics, we examined the association between *z*_PLR_ and Δ*x*_tVNS_. No significant correlation was found across participants (*r* = *−*0.17, *p* = 0.60), suggesting that the additional pupil response to tVNS was independent of natural pupillary light-reflex responsiveness.

#### 3) Reaction-Time Improvement and Stimulation-Induced Pupil Dilation

We examined whether stimulation-induced changes in pupil diameter were associated with changes in visuomotor choice-reaction performance across participants.

For the relationship with the change in gaze time (Δ*T*_1_), the LMM did not find any significant main effect of type (tVNS or Sham) (*b* = 0.041, SE = 0.087, *t*(20) = 0.47, *p* = 0.643), Δ*x*_stim_ (*b* = *−*0.525, SE = 0.608, *t*(20) = *−*0.86, *p* = 0.398), or their interaction (*b* = *−*0.688, SE = 0.737, *t*(20) = *−*0.93, *p* = 0.361) (Fig. 4(i)). Overall, Δ*T*_1_ was not significantly less than zero in tVNS (*t*(11) =*−* 1.61, *p* = 0.137) or Sham (*t*(11) = *−*0.33, *p* = 0.744). These results indicated that gaze time was not reduced with either stimulation, and there was no statistically significant relationship between the change in gaze time and additional pupil response due to either stimulation across participants.

For the relationship with the changes in total reaction time (Δ*T*_2_), there was no main effect of stimulation type (tVNS vs. Sham, with Sham as reference) (*b* = *−*0.021, SE = 0.048, *t*(20) = *−*0.43, *p* = 0.671), whereas an interaction between stimulation type and Δ*x*_stim_ was revealed (*b* = *−*1.721, SE = 0.729, *t*(20) = *−*2.36, *p* = 0.028) (Fig. 4(j)), without a significant main effect of Δ*x*_stim_ (*b* = *−*0.360, SE = 0.683, *t*(20) = *−*0.53, *p* = 0.604). The slope for Δ*T*_2_ as a function of Δ*x*_stim_ was negative and significant in tVNS (slope = *−*2.08 s/mm, 95% CI: [*−* 3.46, *−*0.70], *p* = 0.005) but not significant in Sham (slope = *−*0.36 s/mm, 95% CI: [*−*1.78, 1.06], *p* = 0.604). Overall, Δ*T*_2_ was significantly less than zero in tVNS (*−* 0.133 *±*0.207 s; *t*(11) = *−*2.23, *p* = 0.047), but not in Sham (*−*0.112*±* 0.241 s; *t*(11) = *−*1.62, *p* = 0.134). These findings indicate that reaction time was shorter with tVNS, and participants with greater pupillary responsiveness specific to tVNS (greater Δ*x*_stim_ with tVNS) tend to exhibit a greater reduction in total reaction time (more negative Δ*T*_2_) in a linear manner. In other words, participants with a greater pupil response to tVNS tend to achieve greater performance improvement.

The preceding analyses indicated that stimulation-induced change in total reaction time (Δ*T*_2_) was associated with stimulation-induced pupil dilation for tVNS only (Fig. 4(j)).

In considering extraneous factors, the natural responsiveness of PLR might modulate or confound the stimulation-induced change in total reaction time. We therefore performed an LMM analysis to determine whether these physiological measures are independently associated with the change in total reaction time (Δ*T*_2_) and whether the relationships differed between stimulation types (tVNS vs. Sham). The results of this multi-variate model are reported in Table II.

**TABLE II.** Adjusted Linear Mixed-Effects Model for Δ*T*_2_.

| Predictor | $\beta$ | SE | $t(18)$ | $p$ | 95% CI |
| --- | --- | --- | --- | --- | --- |
| Intercept | -0.648 | 0.532 | -1.22 | 0.238 | [-1.765, 0.469] |
| Type (tVNS vs. Sham) | 0.638 | 0.452 | 1.41 | 0.175 | [-0.311, 1.587] |
| $\Delta x_{\text{stim}}$ | 0.691 | 0.963 | 0.72 | 0.482 | [-1.332, 2.715] |
| $z_{\text{PLR}}$ | 0.014 | 0.014 | 1.01 | 0.324 | [-0.015, 0.042] |
| (tVNS vs. Sham) × $\Delta x_{\text{stim}}$ | -2.469 | 0.893 | -2.77 | **0.013** | [-4.344, -0.593] |
| (tVNS vs. Sham) × $z_{\text{PLR}}$ | -0.017 | 0.012 | -1.47 | 0.160 | [-0.041, 0.007] |
| *Derived simple slopes for $\Delta x_{\text{stim}}$* |  |  |  |  |  |
| Sham | 0.691 | 0.963 | 0.72 | 0.482 | [-1.332, 2.715] |
| tVNS | -1.777 | 0.673 | -2.64 | **0.017** | [-3.191, -0.364] |

Note: $\Delta x_{\text{stim}}$ was centered within stimulation type; Sham was the reference. Simple slopes were derived from the same model. CI: confidence interval; SE: standard error.

When both physiological indices were considered simultaneously (Table II), the interaction between stimulation-induced pupil dilation and stimulation type remained statistically significant, indicating that the relationships between stimulation-induced pupil dilation and the change in reaction time differ between stimulation types. Under tVNS, greater stimulation-induced pupil dilation was associated with greater improvements in total reaction time. While *z*_PLR_ was associated with reaction time in baseline states (Fig. 4(g)), it was not significantly related to the change in total reaction time due to electrical stimulation (Δ*T*_2_). These LMM results confirmed the validity of the findings shown in Fig. 4(j). Collectively, these findings suggest that choice-reaction time during dark transition was associated with natural pupil reactivity, and that reaction time improvement with artificial stimulation depended on stimulation type. Under tVNS, greater stimulation-induced pupil dilation was associated with a greater reduction in reaction time across participants.

## V. Discussion

The major findings of this study are that the transient pupil diameter response to tVNS can be modeled using the LSTM architecture, that the model-optimized tVNS sequence induces pupil dilation in addition to the PLR during the dark transition, and that the tVNS-induced increase in pupil diameter correlates with the improvement of visuomotor performance, i.e., the reduction in choice-reaction time. The leave-one-stimulation-sequence-out cross-validation indicates that the model reliably predicts the pupillary responses for previously unseen stimulation conditions. The observed correlation indicates that individuals with greater tVNS-induced pupil dilation exhibit shorter choice-reaction times during the dark adaptation process. Conversely, the absence of this correlation in the Sham condition suggests that specific tVNS-induced mechanisms behind pupil responses are likely involved in behavioral enhancement.

We hypothesized that tVNS would reduce choice-reaction time through optical and neurophysiological effects associated with enhanced LC–NE activity. Although mean pupil dilation was negatively associated with gaze time (Fig. 4(b)), the tVNS-specific change in pupil dilation was not associated with its reduction (Fig. 4(i)). tVNS increased pupil diameter by 0.11 mm relative to Sham during the first 3 s (Fig. 4(c)), corresponding to an estimated 6.8*±* 5.1% increase in retinal light flux. However, this optical effect may have been too small or variable to improve gaze time. Thus, our findings do not support a meaningful tVNS-induced optical enhancement in healthy young adults. Future studies are warranted to examine whether tVNS could influence this factor if individuals are in a degraded state or population, such as with aging, fatigue, and sleep deprivation.

In contrast, the tVNS-specific change in pupil dilation was significantly associated with the reduction in choice-reaction time (Fig. 4(j)), despite the absence of an association between their mean values in the tVNS session (Fig. 4(d)). Together with the lack of a corresponding association with gaze time, this finding suggests that enhanced LC–NE activity reduces choice-reaction time through a neurophysiological mechanism distinct from the optical effect.

As a second factor, the neurophysiological effect involves multiple neuromodulations and subsequent sensorimotor enhancements. tVNS is regarded to activate the nucleus tractus solitarius and the LC–NE system through the anatomical projections from the auricular branch of the afferent vagus nerve [19], [20]. Additionally, tVNS-induced GABA modulation in certain brain areas [25] may contribute to transient enhancements in cognitive processing [18], motor planning [28], and neuromotor excitability [23]. The currently observed negative correlation between tVNS-induced pupil dilation and the change in choice-reaction time (Fig. 4(j)) may result from accelerated, and possibly fine-tuned, central processing of sensorimotor information [26]. There are studies that reported improvement of reaction time when tVNS was delivered continuously for a prolonged time and without involving a luminance change [25], [27], [28]. The current study advances knowledge by demonstrating that choice-reaction time is improved without continuous prolonged tVNS but with timed brief tVNS, that the improvement is also present during a dark transition, and that the amount of improvement is associated with the magnitude of tVNS-induced pupil dilation across individuals. Since pupil dilation is recognized as reflecting adrenergic activity in the LC [8], [9], this finding implies that individuals with greater neural responsiveness to tVNS may obtain a greater benefit from tVNS.

Our findings demonstrate the technical feasibility of implementing a model-based, feedforward control framework for physiologically adaptive HMIs to mitigate transient operator performance degradation. To ensure timely neuromodulation before sudden illuminance changes (e.g., tunnel entry or surgical lighting fluctuations), the system architecture requires proactive stimulation timing based on the physical latencies identified in this study. Specifically, the developed LSTM network calculates the required tVNS parameters to optimize pupil dilation. Given that the tVNS-induced pupillary response exhibits a latent period of approximately 0.4 s before initiating noticeable dilation under standard conditions, the system must pre-trigger the optimized tVNS sequence at least 0.4 s prior to the predicted luminance transition. Integrating this predictive framework with forward-looking camera streams and foundation models for contextual awareness will enable an automated, feedforward visuomotor enhancement system capable of anticipating dynamic environmental shifts.

The findings in Fig. 4(j) indicate inter-individual variability in tVNS responsiveness; individuals with larger tVNS-induced pupil dilation exhibited greater reductions in choice-reaction time. Although personalized optimization may be feasible, user-specific tuning would require an additional preoperational data-collection phase, increasing setup time before system operation.

This study has several limitations. The sample size (*N* = 12) is relatively modest. Although our findings are supported by consistent results across multiple measures and model-based validation, future studies should aim to confirm these findings with a larger number of participants. Second, the participants were limited to healthy young adults in an unfatigued state without sleep deprivation. The efficacy of the proposed system could be greater and should be evaluated in populations at higher risk of dark adaptation failure, such as elderly, fatigued, or sleep-deprived individuals. A physiological refractory period may exist after tVNS. A temporary desensitization of the LC–NE system has been reported to potentially reduce the efficacy of subsequent stimulation within a short time window [10]. Further investigation is required to evaluate whether this desensitization affects sequential operations, such as driving through successive tunnels. These future studies will advance the understanding and effective utilization of tVNS as an adaptive HMI.

## VI. Conclusions

This study examined whether tVNS can enhance visuomotor reaction performance during the initial phase of dark transition in healthy young adults. The results showed that brief tVNS reliably increased pupil diameter and that this neurophysiological response co-varied with the shortening of the choice-reaction time at the transition from bright to dark environments. We also found that tVNS-induced dilation adds to the PLR even under rapidly changing illumination, and that a tVNS-effect model trained in standard lighting can reproduce comparable responses during dynamic transitions to a dark environment. The analysis of individual differences indicates that participants with greater tVNS-specific responsiveness tend to show greater behavioral improvement. These findings collectively suggest that the model-optimized tVNS approach is highly likely to effectively engage neurophysiological pathways during dark adaptation, offering a promising approach for adaptive HMIs.

## References

[1] G. Q. Yang, T. Chen, Y. Tao, and Z. M. Zhang, “Recent advances in the dark adaptation investigations,” Int. J. Ophthalmol., vol. 8, no. 6, pp. 1245–1252, Dec. 2015.

[2] V. B. Nakagawara, R. W. Montgomery, and K. J. Wood, “Aircraft accidents and incidents associated with visual effects from bright light exposures during low-light flight operations,” Optom. - J. Amer. Optometric Assoc., vol. 78, no. 8, pp. 415–420, Aug. 2007.

[3] D. Zhigang, Z. Zheng, M. Zheng, B. Ran, and X. Zhao, “Drivers’ visual comfort at highway tunnel portals: A quantitative analysis based on visual oscillation,” Transp. Res. Part D, Transp. Environ., vol. 31, pp. 37–47, Aug. 2014.

[4] L. A. Levin, P. L. Kaufman, and M. E. Hartnett, Adler’s Physiology of the Eye, 12th ed. London, U.K.: Elsevier, 2024.

[5] P. Bitsios, R. Prettyman, and E. Szabadi, “Changes in autonomic function with age: A study of pupillary kinetics in healthy young and old people,” Age Ageing, vol. 25, no. 6, pp. 432–438, Nov. 1996.

[6] A. Kaifie, M. Reugels, T. Kraus, and M. Kursawe, “The pupillary light reflex (PLR) as a marker for the ability to work or drive – A feasibility study,” J. Occup. Med. Toxicol., vol. 16, no. 1, Art. no. 43, Nov. 2021.

[7] S. Saito, M. Sotoyama, S. Saito, and S. Taptagaporn, “Physiological indices of visual fatigue due to VDT operation: Pupillary reflexes and accommodative responses,” Ind. Health, vol. 32, no. 2, pp. 57–66, Apr. 1994.

[8] S. Joshi, Y. Li, R. Kalwani, and J. Gold, “Relationships between pupil diameter and neuronal activity in the locus coeruleus, colliculi, and cingulate cortex,” Neuron, vol. 89, no. 1, pp. 221–234, Jan. 2016.

[9] J. Reimer et al., “Pupil fluctuations track rapid changes in adrenergic and cholinergic activity in cortex,” Nat. Commun., vol. 7, Art. no. 13289, Nov. 2016.

[10] G. Aston-Jones and J. D. Cohen, “An integrative theory of locus coeruleus–norepinephrine function: Adaptive gain and optimal performance,” Annu. Rev. Neurosci., vol. 28, pp. 403–450, Jul. 2005.

[11] S. Nieuwenhuis, G. Aston-Jones, and J. D. Cohen, “Decision making, the P3, and the locus coeruleus–norepinephrine system,” Psychol. Bull., vol. 131, no. 4, pp. 510–532, Jul. 2005.

[12] X. Wang et al., “Effects of phasic activation of locus ceruleus on cortical neural activity and auditory discrimination behavior,” J. Neurosci., vol. 44, no. 37, Art. no. e1296232024, Sep. 2024.

[13] C. Rodenkirch, Y. Liu, B. J. Schriver, and Q. Wang, “Locus coeruleus activation enhances thalamic feature selectivity via norepinephrine regulation of intrathalamic circuit dynamics,” Nat. Neurosci., vol. 22, no. 1, pp. 120–133, Jan. 2019.

[14] S. Ghosh and J. H. Maunsell, “Locus coeruleus norepinephrine contributes to visual-spatial attention by selectively enhancing perceptual sensitivity,” Neuron, vol. 112, no. 13, pp. 2231–2240, Jul. 2024.

[15] E. M. Vazey, D. E. Moorman, and G. Aston-Jones, “Phasic locus coeruleus activity regulates cortical encoding of salience information,” Proc. Natl. Acad. Sci. USA, vol. 115, no. 40, pp. E9439–E9448, Oct. 2018.

[16] J. W. de Gee, O. Colizoli, N. A. Kloosterman, T. Knapen, S. Nieuwenhuis, and T. H. Donner, “Dynamic modulation of decision biases by brainstem arousal systems,” eLife, vol. 6, Art. no. e23232, Apr. 2017.

[17] L. Collins, L. Boddington, P. J. Steffan, and D. McCormick, “Vagus nerve stimulation induces widespread cortical and behavioral activation,” Curr. Biol., vol. 31, no. 10, pp. 2088–2098, May 2021.

[18] M. Jigo, J. B. Carmel, Q. Wang, and C. Rodenkirch, “On-demand sensory enhancement via transcutaneous cervical vagus nerve stimulation,” in Brain-Computer Interface Research: A State-of-the-Art Summary 13, C. Guger et al., Eds. Cham, Switzerland: Springer, 2025, ch. 10, pp. 101–112.

[19] T. Kraus et al., “BOLD fMRI deactivation of limbic and temporal brain structures and mood enhancing effect by transcutaneous vagus nerve stimulation,” J. Neural Transm., vol. 114, no. 11, pp. 1485–1493, Nov. 2007.

[20] E. Frangos, J. Ellrich, and B. R. Komisaruk, “Non-invasive access to the vagus nerve central projections via electrical stimulation of the external ear: fMRI evidence in humans,” Brain Stimul., vol. 8, no. 3, pp. 624–636, May/Jun. 2015.

[21] J. W. R. van Leusden, R. Sellaro, and L. S. Colzato, “Transcutaneous vagal nerve stimulation (tVNS): A new neuromodulation tool in healthy humans?” Front. Psychol., vol. 6, Art. no. 102, Feb. 2015.

[22] N. Yakunina, S. S. Kim, and E. Nam, “BOLD fMRI effects of transcutaneous vagus nerve stimulation in patients with chronic tinnitus,” PLOS ONE, vol. 13, no. 11, Art. no. e0207281, Nov. 2018.

[23] C. Perrin et al., “Transcutaneous auricular vagus nerve stimulation during movement selectively activates motor circuitry without additional cortical or autonomic effects,” J. Neurosci., vol. 45, no. 5, Art. no. e2251–24, Feb. 2025.

[24] O. Sharon, F. Fahoum, and Y. Nir, “Transcutaneous vagus nerve stimulation in humans induces pupil dilation and attenuates alpha oscillations,” J. Neurosci., vol. 41, no. 2, pp. 320–330, Jan. 2021.

[25] C. Beste et al., “Effects of concomitant stimulation of the GABAergic and norepinephrine system on inhibitory control – A study using transcutaneous vagus nerve stimulation,” Brain Stimul., vol. 9, no. 6, pp. 811–818, Nov. 2016.

[26] B. J. Jongkees, M. A. Immink, A. Finisguerra, and L. S. Colzato, “Transcutaneous vagus nerve stimulation (tVNS) enhances response selection during sequential action,” Front. Psychol., vol. 9, Art. no. 1159, Jul. 2018.

[27] L. Chen et al., “Effects of transcutaneous vagus nerve stimulation (tVNS) on action planning: A behavioural and EEG study,” IEEE Trans. Neural Syst. Rehabil. Eng., vol. 30, pp. 1675–1683, Jun. 2022.

[28] L. Skora, A. Marzecová, and G. Jocham, “Tonic and phasic transcutaneous auricular vagus nerve stimulation (taVNS) both evoke rapid and transient pupil dilation,” Brain Stimul., vol. 17, no. 2, pp. 233–244, Mar. 2024.

[29] C. Wienke, M. Grueschow, A. Haghikia, and T. Zaehle, “Phasic, event-related transcutaneous auricular vagus nerve stimulation modifies behavioral, pupillary, and low-frequency oscillatory power responses,” J. Neurosci., vol. 43, no. 36, pp. 6306–6319, Sep. 2023.

[30] D. G. Pelli and P. Bex, “Measuring contrast sensitivity,” Vision Res., vol. 90, pp. 10–14, Sep. 2013.

[31] J. Peirce et al., “PsychoPy2: Experiments in behavior made easy,” Behav. Res. Methods, vol. 51, no. 1, pp. 195–203, Feb. 2019.

[32] R. Oostenveld, P. Fries, E. Maris, and J.-M. Schoffelen, “FieldTrip: Open source software for advanced analysis of MEG, EEG, and invasive electrophysiological data,” Comput. Intell. Neurosci., vol. 2011, Art. no. 156869, 2011.

[33] A. Czajka, “Pupil dynamics for iris liveness detection,” IEEE Trans. Inf. Forensics Security, vol. 10, no. 4, pp. 726–735, Apr. 2015.

[34] M. A. Urbin, C. W. Lafe, T. W. Simpson, G. F. Wittenberg, B. Chandrasekaran, and D. J. Weber, “Electrical stimulation of the external ear acutely activates noradrenergic mechanisms in humans,” Brain Stimul., vol. 14, no. 4, pp. 990–1001, Jul./Aug. 2021.

